# GWAS and Fine-Mapping of Livability and Six Disease Traits in Holstein Cattle

**DOI:** 10.1101/775098

**Authors:** Ellen Freebern, Daniel JA Santos, Lingzhao Fang, Jicai Jiang, Kristen L. Parker Gaddis, George E. Liu, Paul M. Vanraden, Christian Maltecca, John B. Cole, Li Ma

**Author notes:** Equal contribution (Ellen Freebern, Daniel JA Santos). Corresponding Author: Li Ma, Department of Animal and Avian Sciences, University of Maryland, College Park, Maryland 20742, USA.

## Abstract

**Background:** Health traits are of significant economic importance to the dairy industry due to their effects on milk production and associated treatment costs. Genome-wide association studies (GWAS) provide a means to identify associated genomic variants and thus reveal insights into the genetic architecture of complex traits and diseases. The objective of this study is to investigate the genetic basis of seven health traits in dairy cattle and to identify potential candidate genes associated with cattle health using GWAS, fine mapping, and analyses of multitissue transcriptome data.

**Results:** We studied cow livability and six direct disease traits, mastitis, ketosis, hypocalcemia, displaced abomasum, metritis, and retained placenta, using de-regressed breeding values and more than three million imputed DNA sequence variants. After data edits and filtering on reliability, phenotypes for 11,880 to 24,699 Holstein bulls were included in the analyses of the seven traits. GWAS was performed using a mixed-model association test, and a Bayesian fine-mapping procedure was conducted to calculate a posterior probability of causality to each variant and gene in the candidate regions. The GWAS results detected a total of eight genome-wide significant associations for three traits, cow livability, ketosis, and hypocalcemia, including the bovine MHC region associated with livability. Our fine-mapping of associated regions reported 20 candidate genes with the highest posterior probabilities of causality for cattle health. Combined with transcriptome data across multiple tissues in cattle, we further exploited these candidate genes to identify specific expression patterns in disease-related tissues and relevant biological explanations such as the expression of *GC* in the liver and association with mastitis as well as the *CCDC88C* expression in CD8 cells and association with cow livability.

**Conclusions:** Collectively, our analyses report six significant associations and 20 candidate genes of cattle health. With the integration of multi-tissue transcriptome data, our results provide useful information for future functional studies and better understanding of the biological relationship between genetics and disease susceptibility in cattle.

## Background

One of the fundamental goals of animal production is to profitably produce nutritious food for humans from healthy animals. Profitability of the dairy industry is influenced by many factors, including production, reproduction, and animal health [1]. Cattle diseases can cause substantial financial losses to producers as the result of decreased productivity, including milk that must be dumped, and increased costs for labor and veterinary care. Indirect costs associated with reduced fertility, reduced production after recovery, and increased risk of culling also can be substantial. For example, ketosis is a metabolic disease that occurs in cows during early lactation and hinders the cow’s energy intake, thus subsequently reduces milk yield and increases the risk of displaced abomasum, which is very costly [2]. Mastitis is a major endemic disease of dairy cattle that can lead to losses to dairy farmers due to contamination, veterinary care, and decreased milk production [3]. In addition, cows may develop milk fever, a metabolic disease that is related to a low blood calcium level known as hypocalcemia [4]. Another common disease in cattle is metritis, which is inflammation of the uterus and commonly seen following calving when cows have a suppressed immune system and are vulnerable to bacterial infection [5]. Complications during delivery can also result in a retained placenta [6]. Despite the complex nature of these cattle diseases, a better understanding of the underlying genetic components can help the management and genetic improvements of cattle health.

Genome-wide association studies (GWAS) have been successful at interrogating the genetic basis of complex traits and diseases in cattle [7–9]. Due to the high level of linkage disequilibrium (LD) between genomic variants, pinpointing causal variants of complex traits has been challenging [9]. Fine-mapping is a common post-GWAS analysis, where posterior probabilities of causality are assigned to candidate variants and genes [9]. In humans, fine-mapping of complex traits are currently on-going along or following GWAS studies [10]. The utility of fine-mapping in cattle studies, however, has been limited by data availability and the high levels of LD present in cattle populations. To circumvent this challenge, a recent study developed a fast Bayesian Fine-MAPping method (BFMAP), which performs fine-mapping by integrating various functional annotation data [9]. Additionally, this method can be exploited to identify biologically meaningful information from candidate genes to enhance the understanding of complex traits [11].

The U.S. dairy industry has been collecting and evaluating economically important traits in dairy cattle since the late 1800s, when the first dairy improvement programs were formed. Since then, a series of dairy traits have been evaluated, including production, body conformation, reproduction, and health traits. Cow livability was included in the national genomic evaluation system by the Council on Dairy Cattle Breeding (CDCB) in 2016 [12]. This trait reflects a cow’s overall ability to stay alive in a milking herd by measuring the percentage of on-farm deaths per lactation. Cow livability is partially attributable to health and can be selected to provide more milk revenue and less replacement of cows. In 2018, six direct health traits were introduced into the U.S. genomic evaluation, including ketosis, mastitis, hypocalcemia or milk fever, metritis, retained placenta, and displaced abomasum [13]. These phenotypic records along with genotype data collected from the U.S. dairy industry provide a unique opportunity to investigate the genetic basis of cattle health. The aim of our study is, therefore, to provide a powerful genetic investigation of seven health traits in cattle, to pinpoint the candidate disease genes and variants with relevant tissue-specific expression, and to provide insights into the biological relationship between candidate genes and the disease risk they may present on a broad scale.

## Results

### Genome-wide association study of livability and six direct health traits

We conducted genome-wide association analyses of seven health related traits in 27,214 Holstein bulls that have many daughter records and thus accurate phenotypes using imputed sequence data and de-regressed breeding values. After editing and filtering on reliability, we included 11,880 to 24,699 Holstein bulls across the seven traits (Table 1). Compared to the analysis using predicted transmitting ability (PTA) as phenotype (Additional File 1), GWAS on de-regressed PTA values produced more consistent and reliable results [14]. While different results between analyses of raw and de-regressed PTAs were obtained for the six health traits, little difference was observed for cow livability, which have more records and higher reliabilities (Table 1 and Additional File 2). Therefore, we only considered association results obtained with de-regressed PTAs in all subsequent analyses.

**Table 1.** Number of Holstein bulls, reliability of PTA, and heritability (h^2^) for six disease traits and cow livability.

| Trait | N | $h^2$ | Average Reliability |
| --- | --- | --- | --- |
| Hypocalcemia | 11,880 | 0.006 | 0.228 |
| Displaced Abomasum | 13,229 | 0.011 | 0.269 |
| Ketosis | 12,468 | 0.012 | 0.260 |
| Mastitis | 14,382 | 0.031 | 0.338 |
| Metritis | 13,653 | 0.014 | 0.281 |
| Retained Placenta | 13,541 | 0.001 | 0.266 |
| Livability | 24,699 | 0.040 | 0.397 |

Out of the seven health traits, we detected significantly associated genomic regions only for three traits after Bonferroni correction, hypocalcemia, ketosis, and livability (Figure 1). In total, we had one associated region on BTA 6 for hypocalcemia, one region on BTA 14 for ketosis, and six regions for cow livability on BTA 5, 6, 14, 18, 21, and 23, respectively (Table 2). Notably, the bovine Major Histocompatibility Complex (MHC) region on BTA 23 [15] is associated with cow livability. Additionally, association signals on BTA 16 for ketosis (*P*-value =1.9×10^-8^) and BTA 6 for mastitis (*P*-value = 4.2×10^-8^) almost reached the Bonferroni significance level. Other traits had prominent signals, but their top associations were below the Bonferroni threshold. Since sequence data have the highest coverage of functional variants in our study, we included all these regions to query the Cattle QTLdb for a comparative analysis.

**Figure 1.**
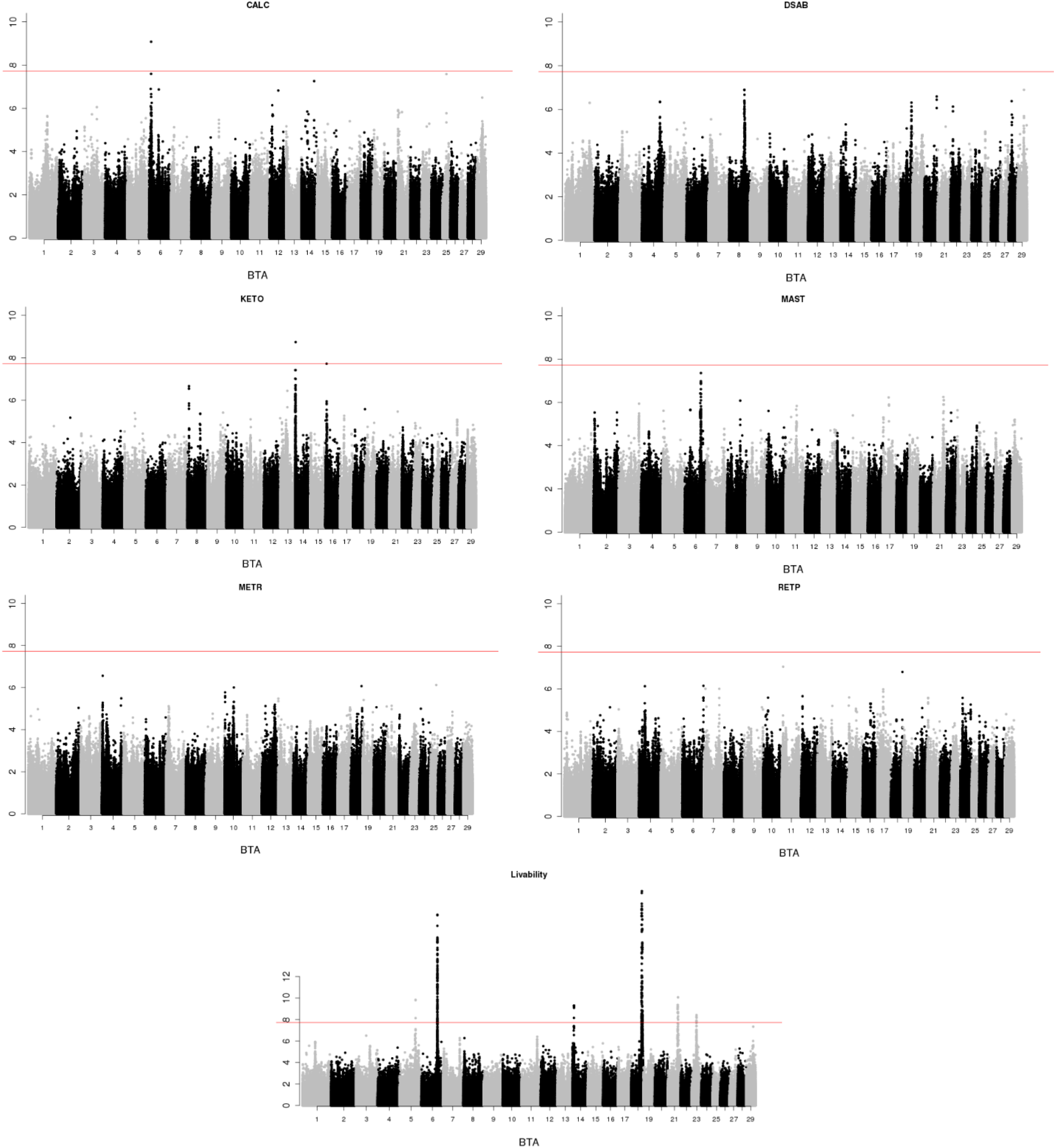
Manhattan plots for hypocalcemia (CALC), displaced abomasum (DSAB), ketosis (KETO), mastitis (MAST), metritis (METR), retained placenta (RETP) and cow livability. The genome-wide threshold (red line) corresponds to the Bonferroni correction.

**Table 2.** Top SNPs and candidate genes associated with hypocalcemia (CALC), displaced abomasum (DSAB), ketosis (KETO), mastitis (MAST), metritis (METR), retained placenta (RETP) and cow livability.

| Trait | Chr | Position | MAF | P-value | Genes Nearby | Traits Previously Associated# |
| --- | --- | --- | --- | --- | --- | --- |
| CALC | 6 | 10,521,824 | 0.014 | $8.3 \times 10^{-10*}$ | *TRAM1L1*,<br>*NDST4* | Subcutaneous fat |
| DSAB | 4 | 97,101,981 | 0.021 | $4.4 \times 10^{-7}$ | *PLXNA4*,<br>*CHCHD3* | Milk protein yield |
| DSAB | 8 | 83,052,202 | 0.109 | $1.3 \times 10^{-7}$ | *FANCC* | Stature |
| DSAB | 29 | 35,977,236 | 0.073 | $1.3 \times 10^{-7}$ | *NTM* | Milk kappa-casein percentage |
| KETO | 14 | 2,762,595 | 0.033 | $1.8 \times 10^{-9*}$ | *LY6K* | Milk protein percentage |
| KETO | 16 | 7,048,452 | 0.019 | $1.9 \times 10^{-8}$ | *KCNT2* | Milk fat percentage |
| MAST | 6 | 88,868,886 | 0.460 | $4.2 \times 10^{-8}$ | *GC* | Clinical mastitis |
| METR | 4 | 3,662,486 | 0.011 | $2.7 \times 10^{-7}$ | *RF00322* | Milk protein yield |
| RETP | 4 | 32,578,298 | 0.218 | $7.4 \times 10^{-7}$ | *RUNDC3B* | Calving ease |
| RETP | 6 | 117,620,548 | 0.026 | $7.2 \times 10^{-7}$ | *QDPR* | Milk kappa-casein percentage |
| RETP | 11 | 7,465,110 | 0.060 | $9.1 \times 10^{-8}$ | *TMEM182* | Abomasum displacement |
| RETP | 18 | 64,492,219 | 0.012 | $1.6 \times 10^{-7}$ | *ZFP28* | Still birth |
| Livability | 5 | 88,823,164 | 0.472 | $1.5 \times 10^{-10*}$ | *ABCC9* | Productive life |
| Livability | 6 | 88,801,999 | 0.454 | $1.7 \times 10^{-18*}$ | *GC* | Clinical mastitis |
| Livability | 14 | 8,536,538 | 0.020 | $5.3 \times 10^{-10*}$ | *ZFAT* | Productive life |
| Livability | 18 | 58,194,319 | 0.075 | $1.1 \times 10^{-20*}$ | *ZNF614* | Bovine respiratory disease |
| Livability | 21 | 56,700,449 | 0.013 | $8.6 \times 10^{-11*}$ | *CCDC88C* | Type |
| Livability | 23 | 26,131,593 | 0.017 | $3.8 \times 10^{-9*}$ | *BLA-DQB* | Antibody-mediated immune response |

\*Genome-wide significance after Bonferroni correction
#Information obtained from the Animal QTLdb for cattle [16]

When compared to existing studies, many of these health related regions have been previously associated with milk production or disease related traits in cattle (Table 2) [16]. The top associated region for hypocalcemia is around 10,521,824 bp on BTA 6, where QTLs were reported for body/carcass weight and reproduction traits with nearby genes being *TRAM1L1* and *NDST4*. The region around 2,762,595 bp on BTA 14 for ketosis is involved with milk and fat metabolism and the well-known *DGAT1* gene. The region around 7,048,452 bp on BTA 16 for ketosis was also previously associated with fat metabolism. The region around 88,868,886 bp on BTA 6 associated with mastitis is close to the *GC* gene with many reported QTLs associated with mastitis [17–20]. This region was also associated with cow livability in this study with QTLs involved with the length of productive life [21]. For the six regions associated with cow livability (Table 2), we found reported QTLs related to productive life, somatic cell count, immune response, reproduction, and body conformation traits [21]. The top associated regions for displaced abomasum (DSAB) on BTA 4 and BTA 8 have been previously associated with cattle reproduction and body conformation traits [22–24]. For metritis, the top associated variant, 3,662,486 bp on BTA4, is close to *RF00322*, and around ±1 Mb upstream and downstream were QTLs associated with production, reproduction, and dystocia [25]. Genes *RUNDC3B* (BTA 4), *QDPR* (BTA 6), *TMEM182* (BTA 11), *ZFP28* (BTA 18) are the closest genes to the retained placenta (RETP) signals with previous associations related to milk production, productive life, health and reproduction traits, including calving ease and stillbirth [8].

### Association of livability QTL with other disease traits

Cow livability is a health-related trait that measures the overall robustness of a cow. As the GWAS of cow livability was the most powerful among the seven traits and detected six QTL regions, we evaluated whether these livability QTLs were also associated with other disease traits. Out of the six livability QTLs, four of them were related to at least one disease trait at the nominal significance level (Table 3). All these overlapped associations exhibited consistent directions of effect: alleles related to longer productive life were more resistant to diseases. The most significant QTL of livability on BTA 18 is associated with displaced abomasum and metritis, both of which can occur after abnormal birth. This QTL has been associated with gestation length, calving traits, and other gestation and birth related traits [11]. The QTL on BTA 6 is associated with hypocalcemia, ketosis, and mastitis. The BTA 21 QTL is associated with hypocalcemia and mastitis. The BTA 5 QTL is related to displaced abomasum and ketosis. Interestingly, the bovine MHC region on BTA 23 is not associated with any direct disease traits, which suggests that those genes do not explain substantial variation for the presence or absence of a disease during a lactation.

**Table 3.** Association results of the top SNPs associated with cow livability for hypocalcemia, displaced abomasum (DSAB), ketosis, mastitis, and metritis. *P*-values larger than 0.05 and their Beta coefficients were excluded.

| Chr | Position | Livability |  | Hypocalcemia |  | DSAB |  | Ketosis |  | Mastitis |  | Metritis |  |
| --- | --- | --- | --- | --- | --- | --- | --- | --- | --- | --- | --- | --- | --- |
|  |  | *P*-value | Beta | *P*-value | Beta | *P*-value | Beta | *P*-value | Beta | *P*-value | Beta | *P*-value | Beta |
| 5 | 88,823,164 | $1.5 \times 10^{-10}$ | -0.43 | - | - | 0.04 | -0.14 | 0.04 | -0.21 | - | - | - | - |
| 6 | 88,801,999 | $1.7 \times 10^{-18}$ | -0.66 | $5.0 \times 10^{-3}$ | -0.2 | - | - | $2.1 \times 10^{-3}$ | -0.35 | $4.2 \times 10^{-7}$ | -0.75 | - | - |
| 14 | 8,536,538 | $5.3 \times 10^{-10}$ | -1.1 | - | - | - | - | - | - | - | - | - | - |
| 18 | 58,194,319 | $1.1 \times 10^{-20}$ | -1.0 | - | - | $1.1 \times 10^{-4}$ | -0.47 | - | - | - | - | 0.01 | -0.51 |
| 21 | 56,700,449 | $8.6 \times 10^{-11}$ | -1.5 | 0.03 | -0.58 | - | - | - | - | $9.1 \times 10^{-3}$ | -1.43 | - | - |
| 23 | 26,131,593 | $3.8 \times 10^{-9}$ | 0.71 | - | - | - | - | - | - | - | - | - | - |

### Fine-mapping analyses and validation from tissue-specific expression

Focusing on the candidate QTL regions in Table 2, the fine-mapping analysis calculated posterior probabilities of causalities (PPC) for individual variants and genes to identify candidates (Table 4), which were largely consistent with the GWAS results. A total of eight genes detected in GWAS signals were also successfully fine-mapped, including *PLXNA4* (DSAB), *FANCC* (DSAB), *NTM* (DSAB), *GC* (mastitis and livability), *ABCC9* (livability), *QDPR* (RETP), *ZFAT* (livability), and *CCDC88C* (livability). In addition, fine-mapping identified new candidate genes, including *COBL* on BTA 4 for metritis, *LOC783947* on BTA 16 for ketosis, *LOC783493* on BTA 18 for RETP, and *LOC618463* on BTA 18 and *LOC101908667* on BTA 23 for livability. The genes *LOC107133096* on BTA 14 and *LOC100296627* on BTA 4 detected respectively for ketosis and RETP by fine mapping were close to two genes (*DGAT1* and *ABCB1*) that have known biological association with milk production and other traits. In addition to the detected genes in these two cases, we further investigated genes with a potential biological link with disease, and genes with the highest PPC (*PARP10* and *MALSU1*) that were located between these two references (Table 4). No genes were detected by fine-mapping in the signal on BTA 6 for hypocalcemia (Figure 1), given that the nearest genes were beyond a 1 Mb window boundary.

**Table 4.**
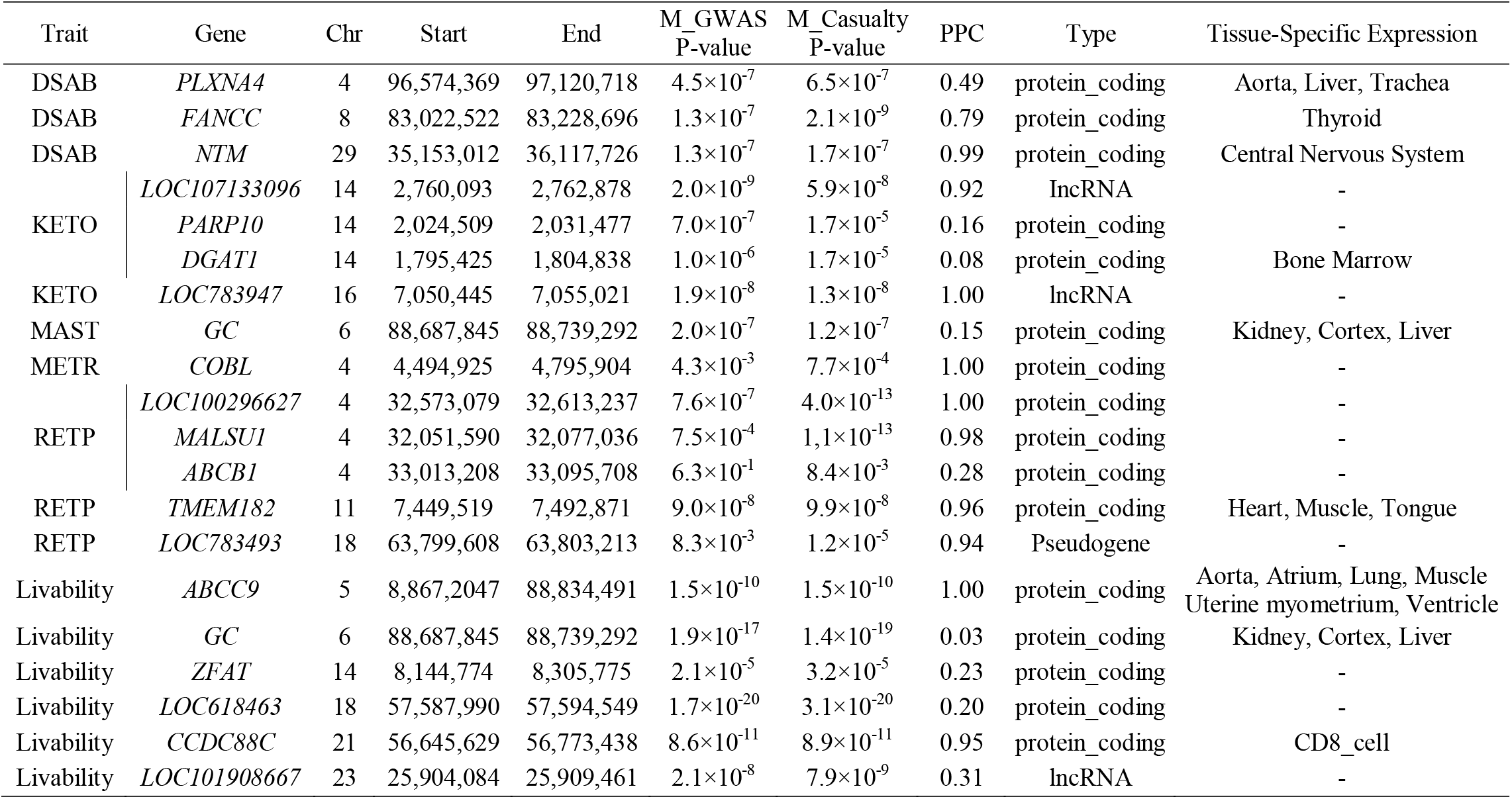
List of candidate genes with highest posterior probability of causality (PPC) and their minimum *P*-values for casualty (M_Causality) and GWAS (M_GWAS) associated with hypocalcemia (CALC), displaced abomasum (DSAB), ketosis (KETO), mastitis (MAST), metritis (METR), retained placenta (RETP) and cow livability and their tissue specific expression.

In addition, we investigated the expression levels of fine-mapped candidate genes across cattle tissues using existing RNA-Seq data from public databases. While many genes are ubiquitously expressed in multiple tissues, several fine-mapped genes were specifically expressed in a few tissues relevant to cattle health (Table 4). Interesting examples of tissuespecific expression and candidate genes included liver with mastitis and livability (*GC*), and CD8 cells with livability (*CCDC88C*). Although this analysis is preliminary, these results provide additional support for these candidate genes of cattle health and help the understanding of how and where their expression is related with dairy disease resistance.

## Discussion

In this study, we performed powerful GWAS analyses of seven health and related traits in Holstein bulls. The resulting GWAS signals were further investigated by a Bayesian fine-mapping approach to identify candidate genes and variants. Additionally, we included tissuespecific expression data of candidate genes to reveal a potential biological relationship between genes, tissues and cattle diseases. Finally, we provide a list of candidate genes of cattle health with associated tissue-specific expression that can be readily tested in future functional validation studies.

In our GWAS analysis, we used de-regressed PTA as phenotype and incorporated the reliabilities of the de-regressed PTAs of livability and six disease traits. Three traits were found to have significant association signals, hypocalcemia, ketosis, and livability, which demonstrated the power of our GWAS study. For instance, we observed tissue-specific expression with hypocalcemia for body/carcass weight and reproduction traits that are supported by a previous study where hypocalcemia compromised immunosuppression in cows at calving [26]. Interestingly, this finding coincides with the incidence of hypocalcemia that occurs in human infants born to mothers who had complications during pregnancy or labor [27]. This suggests that the association signals related to hypocalcemia in these regions may be a common phenomenon across species. In addition to hypocalcemia, we also observed regions associated with livability, in particular, with the region around 58,194,319 on BTA 18 to possess a large effect on dairy and body traits. Our finding was corroborated by a BLAST analysis that identified a related molecule, Siglec-6, which is expressed in tissues such as the human placenta [28]. Further analyses can be performed to characterize the functional implications of these association regions for the seven health and related traits in cattle.

When using PTA values as phenotype in GWAS, we observed different regions to be associated, compared to the GWAS with de-regressed PTA (Figure 1 and Additional File 2). For example, a genomic region larger than 4 Mb on BTA 12 was associated with most of the health traits (Additional File 2). Although these generally appeared as clear association signals, we observed only a few HD SNP markers to be associated, which may be due to poor imputation. Additionally, this region was reported by VanRaden et al., (2017) as having low imputation accuracy [29]. The lower imputation accuracy on BTA 12 was determined to be caused by a gap between the 72.4 and 75.2 Mb region where no SNPs were present on the HD SNP array [29]. Additional studies are needed to address this imputation issue in order to improve the accuracy and power of future analysis on this region. In sum, this comparison of GWAS using PTA and de-regressed PTA supports the use of de-regressed PTA values with reliabilities accounted for in future GWAS studies in cattle.

Application of BFMAP for fine-mapping allowed us to identify 20 promising candidate genes (Table 4) and a list of candidate variants (Additional File 3) for health traits in dairy cattle. We found that most of the genes possess tissue-specific expression, notably the detected gene *LOC107133096* on BTA 14 for ketosis. This gene is located close to the *DGAT1* gene that affects milk fat composition. A previous candidate gene association study by Tetens et al. (2013) proposed *DGAT1* to be an indicator of ketosis [30]. In that study, the *DGAT1* gene was determined to be involved in cholesterol metabolism, which is known to be an indicator of a ketogenic diet in humans [30]. This result highlights a potential pathway in the pathogenesis of ketosis that may be an area for future research. Additionally, ketosis is a multifactorial disease that is likely influenced by multiple loci. Therefore, implementation of a functional genomics approach would allow identification of more genetic markers, and in doing so, improve resistance to this disease. For DSAB, the gene *PLXNA4* was observed to have an association with the variant 97,101,981 bp on BTA 4 (Table 4 and Additional File 3). Our analysis also detected tissue-specific expression for *PLXNA4* in the aorta. A previous study on atherosclerosis found that Plexin-A4 knockout mice exhibited incomplete aortic septation [31]. In addition, it is worth noting that one of the known clinical signs of dairy cows with displaced abomasum is an increased heart rate. Taken together, these findings provide support for the association of *PLXNA4* with DSAB as we found in this study.

Six signals were observed as clear association peaks for livability (Figure 1). The associated variant at 8,144,774 – 8,305,775 bp on BTA 14 was close to the gene *ZFAT*, which is known to be expressed in the human placenta [32]. In particular, the expression of this gene is downregulated in placentas from complicated pregnancies. Additionally, a GWAS study performed in three French dairy cattle populations found the *ZFAT* gene to be the top variant associated with fertility [33]. These results lend support to our findings of this associated variant with the livability health trait. On BTA18, the associated variant at 57,587,990 – 57,594,549 bp was near the gene *LOC618463*, which has been previously identified as a candidate gene associated with calving difficulty in three different dairy populations [34]. For the associated variant at 56,645,629 – 56,773,438 bp on BTA21, it is located close to the *CCDC88C* gene (Table 4). In addition to our detection of tissue-specific expression with the CD8 cell, this gene has been associated with traits such as dairy form and days to first breeding in cattle [19]. Interestingly, in humans, mutation of the *CCDC88C* gene caused a recessive form of congenital hydrocephalus, which presents symptoms of an elevated CD4/CD8 ratio [35, 36].

It is notable that our GWAS signal for livability at 25,904,084 – 25,909,461 bp on BTA 23 is located in the bovine MHC region (Table 4). The gene we detected was *LOC101908667*, which is one of the immune genes of MHC. This is of considerable interest because MHC genes have a role in immune regulation. The MHC complex of cattle located on BTA 23 is called the bovine leukocyte antigen (*BoLA*) region. This complex of genes has been extensively studied, such as in research investigating the polymorphism of genes in *BoLA* and their association with disease resistance [37]. Therefore, our research highlights a gene of considerable interest that should be further explored to understand its importance in breeding programs and its potential role in resistance to infectious diseases.

Additionally, we identified an associated variant for livability at 88,687,845 - 88,739,292 bp on BTA6 was close to the gene *GC*, which was specifically expressed in tissues such as the liver (Table 4). This gene has been previously studied in an association analysis that investigated the role of *GC* on milk production [17]. It found that the gene expression of *GC* in cattle is predominantly expressed in the liver. Moreover, affected animals displayed decreased levels of the vitamin D-binding protein (DBP) encoded by *GC*, highlighting the importance of *GC* for a cow’s production. Additionally, liver-specific *GC* expression has been identified in humans, specifically regulated through binding sites for the liver-specific factor HNF1 [38]. Collectively, these results offer evidence for *GC* expression in the liver, which may be an important factor for determining cow livability.

Interestingly, the *GC* gene was also detected to have tissue-specific expression in the liver for mastitis (Table 4). This is corroborated by a study on cattle infected with mastitis to possess limited DBP concentration [17]. Vitamin D plays a key part in maintaining serum levels of calcium when it is secreted into the milk [39]. Since *GC* encodes DBP, it was suggested that the *GC* gene has a role in regulating milk production and the incidence of mastitis infection in dairy cattle. It is important to note that bovine mastitis pathogens, such as *Staphylococcus aureus* and *Escherichia coli*, also commonly occur as pathogens of humans. Therefore, development of molecular methods to contain these pathogens is of considerable interest for use in human medicine to prevent the spread of illness and disease. For instance, the use of enterobacterial repetitive intergenic consensus typing enables trace back of clinical episodes of *E. coli* mastitis, thus allowing for an evaluation of antimicrobial products for the prevention of mastitis [40]. Continued investigation using molecular methods are needed to understand the pathogenesis of mastitis and its comparative relevance to human medicine. Based on the fine mapping for metritis, the new gene assigned was *COBL* on BTA 6 (Table 4). However, this candidate gene was found to have variants only passing the nominal significance level for causality and for GWAS. Further exploration of this candidate gene is needed to contribute to our understanding of its function and potential tissue-specific expression.

For RETP, the gene *TMEM182* was observed to have an association with a variant between 7,449,519 – 7,492,871 bp on BTA11 (Table 4). Our tissue-specific analysis identified *TMEM182* to have an association in muscle tissues. A study performed in Canchim beef cattle investigated genes for male and female reproductive traits and identified *TMEM182* on BTA 11 as a candidate gene that could act on fertility [41]. Additionally, the gene *TMEM182* has been found to be up-regulated in brown adipose tissue in mice during adipogenesis, which suggests a role in the development of muscle tissue [42]. One important factor that causes retention of fetal membranes in cattle is the impaired muscular tone of organs such as the uterus and abdomen [43]. This suggests the importance of the *TMEM182* gene and the need for future studies to better understand its role in the cattle breeding program.

## Conclusions

In this study, we reported eight significant associations for seven health and related traits in dairy cattle. In total, we identified 20 candidate genes of cattle health with the highest posterior probability, which are readily testable in future functional studies. Several candidate genes exhibited tissue-specific expression related to immune function, muscle growth and development, and neurological pathways. The identification of a novel association for cow livability in the bovine MHC region also represented an insight into the biology of disease resistance. Overall, our study offers a promising resource of candidate genes associated with complex diseases in cattle that can be applied to breeding programs and future studies of disease genes for clinical utility.

## Methods

### Ethics statement

This study didn’t require the approval of the ethics committee, as no biological materials were collected.

### Genotype data

Using 444 ancestor Holstein bulls from the 1000 Bull Genomes Project as reference, we previously imputed sequence variants for 27,214 progeny-tested Holstein bulls that have highly reliable phenotypes via FindHap version 3 [44]. The original 777,962 HD SNPs were reduced to 312,614 by removing highly correlated SNP markers with a |r| value higher than 0.95 and by prior editing. Variants with a minor allelic frequency (MAF) lower than 0.01, incorrect map locations (UMD3.1 bovine reference assembly), an excess of heterozygotes, or low correlations (|r| < 0.95) between sequence and HD genotypes for the same variant were removed. The final imputed data was composed of 3,148,506 sequence variants for 27,214 Holstein bulls. Details about the genomic data and imputation procedure are described by VanRaden et al., (2017). After imputation, we only retained autosomal variants with MAF ≥0.01 and *P*-value of Hardy-Weinberg equilibrium test >10^-6^.

### Phenotype data

The data used were part of the 2018 U.S. genomic evaluations from the Council on Dairy Cattle Breeding (CDCB), consisting of 1,922,996 Holstein cattle from the national dairy cattle database. De-regressed predicted transmitting ability (PTA) values according to Garrick et al., (2009) for livability, hypocalcemia, displaced abomasum, ketosis, mastitis, metritis, and retained placenta were analyzed. We restricted the de-regression procedure to those bulls with PTA reliability greater than parent average reliability, thus reducing the total number of animals from 27,214 to 11,880, 13,229, 12,468, 14,382, 13,653, 13,541, and 24,699 for the seven traits, respectively (Table 1).

### Genome-wide association study (GWAS)

A mixed model GWAS approach was performed using MMAP [45]. Thus, the additive effect was divided into a random polygenic effect and a fixed effect of the candidate SNP. The variance components were estimated using REML. The model can be generally presented as:

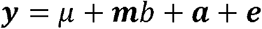

where ***y*** is a vector with de-regressed PTAs; ***μ*** is the global mean; ***m*** is the candidate SNP genotype (allelic dosage coded as 0, 1 or 2) for each animal; ***b*** is the solution effect of the candidate SNP; ***a*** is a solution vector of polygenic effect accounting for the population structure assuming 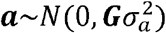, where **G** is a relationship matrix; and ***e*** is a vector of residuals assuming 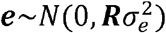, where **R** is a diagonal matrix with diagonal elements weighted by the individual de-regressed reliability 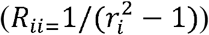. For each candidate variant, a Wald test was applied to evaluate the alternative hypothesis, H_1_: *b* ≠ 0, against the null hypothesis H_0_: *b* = 0. Bonferroni correction for multiple comparisons was applied to control the type-I error rate. Gene coordinates in the UMD v3.1 assembly [46] were obtained from the *Ensembl Genes 90* database using the *BioMart tool* [47]. The *cattle QTLdb* database [16] was examined to check if any associated genomic region was previously reported as a cattle quantitative trait locus (QTL).

### Fine-mapping association study

In order to identify potential candidate genes and their causal variants, GWAS signals were investigated through a fine-mapping procedure using a Bayesian approach with the software BFMAP v.1 [19]. Independent association signals within the same QTL region as well as a posterior probability of causality (PPC) to each variant and its causality *p-value* were assigned. The minimal region covered by each lead variant was determined as ±1 Mb upstream and downstream (candidate region ≥2 Mb). This extension allowed the region to cover most variants that have an LD r^2^ of >0.3 with the lead variants. The employed fine-mapping approach included three steps: forward selection to add independent signals in the additive Bayesian model, repositioning signals, and generating credible variant sets for each signal. Details about the BFMAP algorithm and its procedure are described by Jiang et al., (2019).

### Tissue-specific expression of candidate genes

We have collected and analyzed RNA-seq data of 723 samples from 91 tissues and cell types in Holstein cattle from previous studies on NCBI GEO database. Details of the processing and analyses have been described in a submitted manuscript [48]. We processed all the 732 RNA-seq data uniformly using a rigorous bioinformatics pipeline with stringent quality control procedures. We then calculated the *t*-statistics for differential expression for each gene in each tissue in the RNA-seq dataset using a previous method [49]. Specifically, the log2-transformed expression (i.e., log2FPKM) of genes was standardized with mean of 0 and variance of 1 within each tissue or cell type,

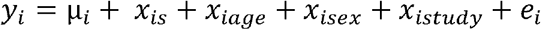

where *y_i_* is the standardized log2-transformed expression level (i.e., log2FPKM) of *i*th gene; μ_*i*_ is the overall mean of the *i*th gene; *x_is_* is the tissue effect, where samples of the tested tissue were denoted as ‘1’, while other samples as ‘-1’; *x_iage_, x_isex_, x_istudy_* were age, sex, and study effects for the *i*th gene, respectively; *e_i_* is residual effect. We fitted this model for each gene in each tissue using the ordinary least-square approach and then obtained the t-statistics for the tissue effect to measure the expression specificity of this gene in the corresponding tissue. Using this approach, we evaluated the expression levels for each of the candidate genes that were fine-mapped in this study across the 91 tissues and cell types and identified the most relevant tissue or cell type for a disease trait of interest.

## List of abbreviations

GWAS: genome-wide association study.
HO: Holstein.
SNP: single nucleotide polymorphism.
QTL: quantitative trait locus.
BTA: Bos taurus chromosome.
CALC: hypocalcemia.
LD: Linkage disequilibrium.
MAF: minor allelic frequency.
PPC: probability of causality.
PTA: Predicted Transmitting Ability.
CALC: hypocalcemia.
DSAB: displaced abomasum.
KETO: Ketosis.
MAST: Mastitis.
METR: Metritis.
RETP: retained placenta.

## Declarations

### Ethics approval and consent to participate

The Council on Dairy Cattle Breeding (CDCB) approved data access and the research, which was conducted on USDA-AGIL computers using the national database shared jointly by CDCB and AGIL under a non-funded cooperative agreement: http://aipl.arsusda.gov/reference/CDCB_NFCA.pdf.

### Consent for publication

Not applicable.

### Availability of data and material

The original performance and pedigree data are owned by CDCB. A request to CDCB to access the data may be sent to: João Dürr, CDCB Chief Executive Officer. Bull genotypes are controlled by the Collaborative Dairy DNA Repository (CDDR; Verona, WI), and a request to access those data must be made to Jay Weiker, CDDR Administrator. The bovine transcriptome data can be directly downloaded from NCBI GEO database with accession numbers SRP042639, PRJNA177791, PRJNA379574, PRJNA416150, PRJNA305942, SRP111067, PRJNA392196, PRJNA428884, PRJNA298914, PRJEB27455, PRJNA268096, and PRJNA446068. All other data and results are included in the published article.

### Authors’ contributions

LM, JC, and CM conceived the study. EF, DS, LF, JJ analyzed and interpreted data. EF, DS, CM and LM wrote the manuscript. GEL, KPG, JC, PMV contributed tools and materials. All authors read and approved the final manuscript.

## Acknowledgements

We thank the 1000 Bull Genomes Project for providing reference sequence data for imputation. The Council on Dairy Cattle Breeding (CDCB; Bowie, MD), Cooperative Dairy DNA Repository (Verona, WI), and dairy industry contributors are thanked for providing phenotypic, pedigree, and genomic data. Mention of trade names or commercial products in this article is solely for the purpose of providing specific information and does not imply recommendation or endorsement by the US Department of Agriculture. The USDA is an equal opportunity provider and employer.

## Additional Files

**Additional File 1:**
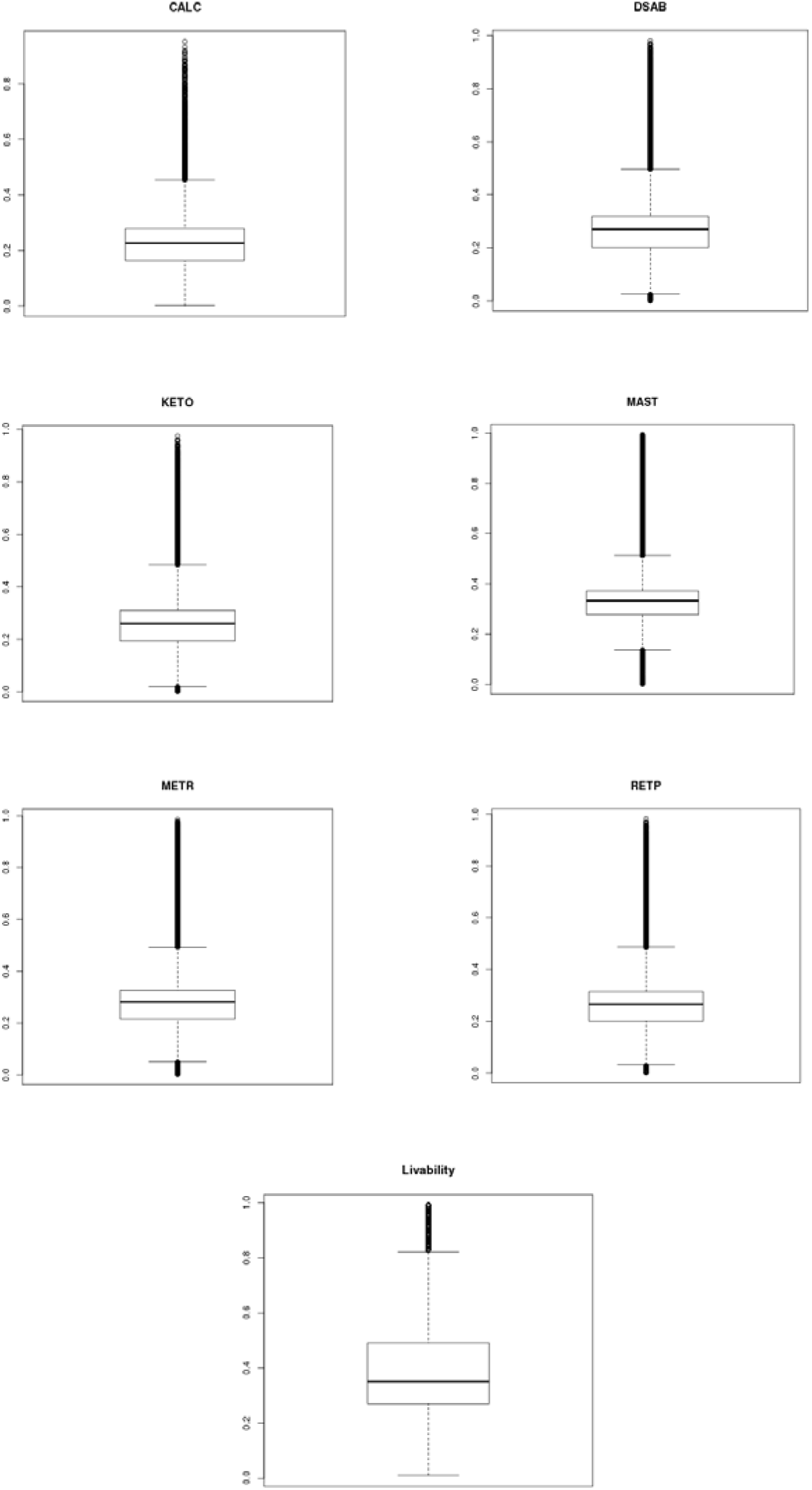
Boxplot with PTA reliability for hypocalcemia (CALC), displaced abomasum (DSAB), ketosis (KETO), mastitis (MAST), metritis (METR), retained placenta (RETP) and cow livability.

**Additional File 2:**
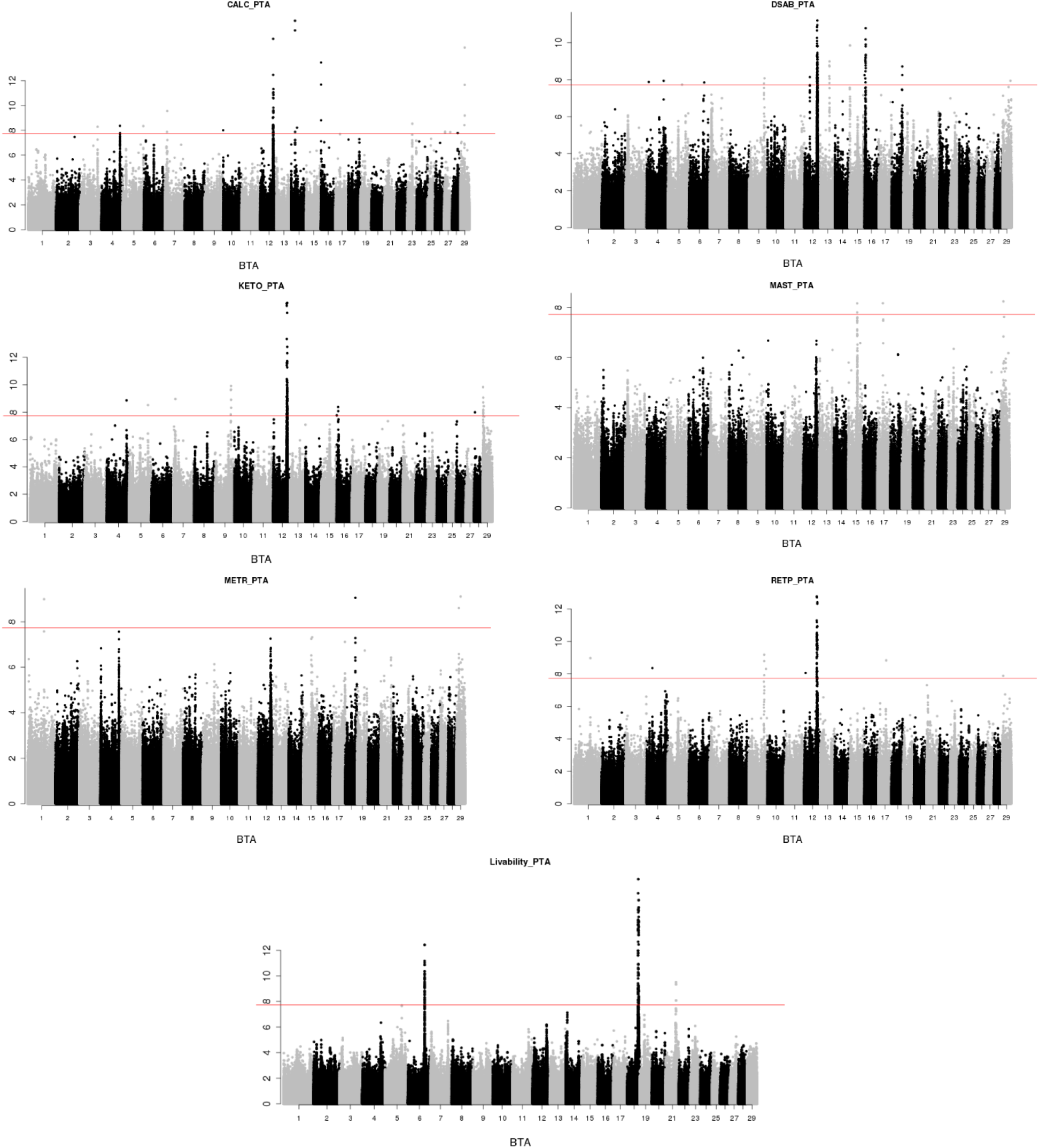
Manhattan plots using the PTA as phenotype for hypocalcemia (CALC), displaced abomasum (DSAB), ketosis (KETO), mastitis (MAST), metritis (METR), retained placenta (RETP) and cow livability. The genome-wide threshold (red line) corresponds to the Bonferroni correction for a nominal *P*-value□=□0.05.

**Additional File 3:**
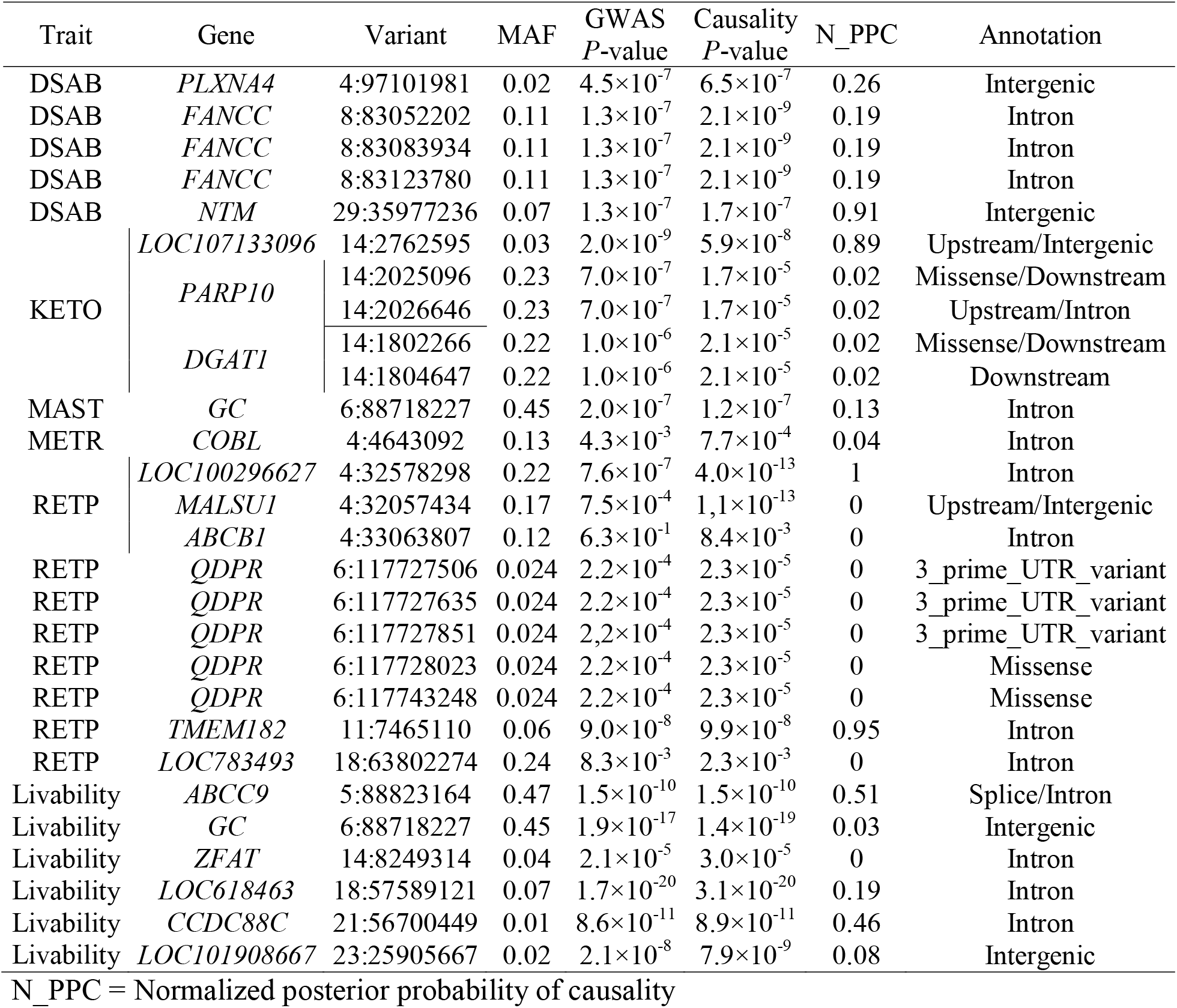
List of variants into genes with highest posterior probability of causality mostly associated with displaced abomasum (DSAB), ketosis (KETO), mastitis (MAST), metritis (METR), retained placenta (RETP) and cow livability.

